# Object-based attentional selection emerges late in visual cortical hierarchy for objects of varying perceptual strengths

**DOI:** 10.1101/2020.05.10.087544

**Authors:** Shahd Al-Janabi, Nofar Strommer, Shai Gabay, Adam S. Greenberg

**Affiliations:** Department of Psychology, University of Wisconsin – Milwaukee, Milwaukee, WI, USA; Department of Psychology, University of Haifa, Haifa, Israel; The Institute of Information Processing and Decision Making (IIPDM), Israel; Department of Biomedical Engineering, Medical College of Wisconsin & Marquette University, Milwaukee, WI, USA

**Author notes:** Corresponding author: Adam S. Greenberg, PhD, Department of Biomedical Engineering, Medical College of Wisconsin & Marquette University, Milwaukee, WI, USA 53226.

## Abstract

Object-based attention (OBA) can - in addition to acting upon explicit object representations - act upon occluded and illusory objects. It remains unknown, however, whether or not the selection of such object representations is detectable at the same level within visual cortex. This study examined the level within visual cortex (V1-V3, LOC) at which object-based selection is observed for explicit, occluded and illusory objects. During fMRI acquisition, participants identified a target preceded by a predictive arrow cue in the double-rectangle cueing paradigm. We independently localized retinotopically-specific regions of cortex corresponding to all possible target locations to examine neural fluctuations at each level of the visual cortical hierarchy. We found, after cue onset, that activity along visual cortex was not greater for representations of cued than of uncued same object locations. In contrast, we found that activity in V3 was enhanced at retinotopic representations that correspond to uncued same than different object locations. These results, together, support attentional spreading. Additionally, when the target appeared at either the cued or uncued locations, we found higher activation in areas representing uncued same object versus cued locations. This effect emerged along the visual cortical hierarchy. Further, when the target appeared on either the cued or uncued object, we found that activation in V3 transiently increased at uncued same than different object locations. This effect was also detectable upstream in LOC. These results index attentional re-orienting between locations/objects. Effects emerged regardless of object type: explicit or completed. Thus, the gating of object information proceeds completion.

**Significance Statement:** We investigated the level within visual cortex (V1-V3, LOC) that object-based selection is observed for explicit objects and those requiring perceptual completion. We showed that activity along visual cortex was similar for representations of locations on a cued object, which may indicate attention spreads evenly to all locations on an object marked as relevant by the cue. We also showed that activity in late visual areas was greater for representations of uncued same than different object locations, which may indicate that attention enhances the cued object. These findings support the attentional spreading account. Object selection may, thus, be instantiated by even engagement of locations within a cued, and/or suppression of locations within an uncued, object - independent of its type.

---

The way in which we perceive our world relies on both the information that enter our senses, and upon which aspects of the information we select. Visual attention is the mechanism by which this selection is conducted, and our ability to navigate seamlessly in our visually cluttered environment is attributed to both space- and object-based selection. These two attentional domains often work in concert, as studies using the double-rectangle cueing paradigm show (Egly, Driver & Rafal, 1994). In this paradigm, participants are presented with two rectangles, followed by a brief illumination of one rectangle’s end (cue), then the filling-in of one rectangle’s end (target). The target appears at the valid location, the invalid location within the same rectangle, or the equidistant (from the cue) invalid location within the different rectangle. Participants respond faster to valid than invalid targets. This *cue validity effect* indexes cued location selection. Participants also respond faster to targets appearing at the invalid same-than different-object location. This *same-object advantage* indexes cued object selection. Interestingly, with respect to this latter phenomenon, the same-object advantage emerges not only on object representations that are explicit, but also on those that require perceptual completion (occluded and illusory rectangles; Moore, Yantis & Vaughan, 1998). This finding was taken to indicate that object-based attention follows amodal and modal completion, which possibly commences early in visual information processing. Consequently, a same-object advantage, which does not vary as a function of object explicitness, is yielded. An implication of this ‘completion *then* attention’ account is that object-based effects should emerge at the same visual cortical hierarchy level for object representations that require perceptual completion versus not. We tested this prediction.

Previous studies have shown that object-based effects produced by explicit objects are observable in visual cortex. Müller and Kleinschmidt (2003), for instance, asked participants to complete a double-rectangle paradigm. In their study, an arrow cue directed attention to the upper left location of explicit objects. The target was 70% likely to appear at that location. Following cue onset, a pattern of BOLD signal enhancement that was greatest for retinotopic representations of the cued object, followed by uncued same object then uncued different object locations emerged, which suggests attentional selection of retinotopic representations follows an object outline, and is organized as a gradient. Critically, this descending activity pattern was present along V1-V4, thus indicating that explicit object-based information is similarly gated along the visual hierarchy. These findings were extended upon by Shomstein and Behrmann (2006) who asked participants to hold attention at a given location within an explicit rectangle, or shift attention to another location within the same/different rectangle, depending on target colour. Following target onset, the researchers found that activation in each retinotopic representation of the target location followed the same graded pattern as reported by Müller and Kleinschmidt (2003), wherein its velocity was greatest when attention was held at a given location, followed by when a shift within an object was elicited then when a shift between an object was elicited. In contrast to Müller and Kleinschmidt (2003), this pattern grew along V1–4, as opposed to remaining constant. Nonetheless, these results, together, suggest object-based effects for explicit objects emerge along visual cortex.

Our primary aim was to examine the locus of object-based effects along the visual cortical hierarchy (V1-V3, LOC) for object representations that do (occluded, illusory) or do not (explicit) require perceptual completion, using functional magnetic resonance imaging (fMRI) with blood oxygenation level-dependent (BOLD) contrast. We measured changes in visual activity at retinotopic representations of cued and uncued (same/different) locations after cue and target onset. We predicted that all objects would yield object-based effects in visual cortex, given that Moore et al. (1998) show they produce an indistinguishable behavioral same-object advantage. But, the extent to which these object-based effects emerge at the same level of visual cortex may depend on whether or not perceptual completion is engaged before attention selects an object.

## Materials and Methods

### Participants

Twenty-two participants (11 females, 11 males; mean + SD age, 27 + 8 years) at the University of Wisconsin–Milwaukee (UWM) participated in the study. All were monetarily reimbursed for participation. The UWM Institutional Review Board approved the study, and it was conducted in conformity with the Declaration of Helsinki.

### Experimental Task

#### Stimuli

Three object types were used in this experiment: **(1)** explicit objects, wherein a pair of gray rectangles measuring 2.69 x 15.01^°^ of visual angle were presented either horizontally or vertically; **(2)** occluded objects, wherein the same explicit rectangles, in addition to an occluding rectangle, were presented. The occluding rectangle measured 17.95^°^ x 2.69^°^ of visual angle, and was oriented orthogonally to the rectangular pair. Moreover, the occluding rectangle was presented stereoscopically in front of the rectangular pair, such that the rectangular pair had to be perceptually completed; and, **(3)** illusory objects, wherein gray circles that subtended 3.20^°^ of visual angle in diameter were presented. The rectangular cutouts in the gray circles, which served as inducing disks, were the same size as the explicit objects, but appeared in black. The cutouts were presented within the circles: 1.60^°^ from the horizontal edge, and 0.27^°^ from the vertical edge. Thus, the resulting perceptual rectangles were equivalent to those presented in the explicit and occluded object conditions. The object pair – in all 3 conditions - were positioned 7.64^°^ apart (edge-to-edge), and a white fixation cross (20-point Monaco font) was centered between them. A white arrow (<) cue was used. The cue was displayed in 20-point Monaco font, and was angled 45^°^ to the upper left, upper right, lower left or lower right. Targets were either the letters X or O, whereas the distractors were the numerical character 4. All target and distractor stimuli were white and appeared in 20-point Monaco font. The targets and distractors appeared 0.82^°^ from the horizontal edge of the rectangles, and 0.63^°^ from the vertical edge of the rectangles. Stimuli were presented on a black background using a custom-made screen, which measured 29.8 cm x 6.7 cm, attached to a NOVA-32 channel head coil. The projection resolution to the screen was 1024 x 768. Participants were positioned horizontally 20.32 cm away from the screen.

#### Experimental Design

fMRI data were acquired over 2 sessions (2 h each). The manipulated within-subjects factors were Object Type (explicit, occluded, illusory), Object Orientation (horizontal, vertical), and Cue-Target Validity (valid, invalid same, invalid different). Object type and orientation were varied randomly within participants, across runs. Three runs of each object type were completed during each session. Each run contained 40 trials and lasted 360 s. Thus, participants completed 18 total functional runs of the experimental task. We also acquired 2 runs of 3 localizer tasks for independent selection of ROIs. Stimulus presentation was controlled by a laptop using the Psychtoolbox-3 extensions (Brainard, 1997; Pelli, 1997; Kleiner et al, 2007) in GNU Octave (Eaton, Bateman, Hauberg, Wehbring, 2015), which was triggered by the scanner.

#### Behavioral procedure

Each run began and ended with a 20 s fixation display. The block structure was as follows (Fig. 1): A central fixation cross was presented along with two rectangles oriented either horizontally or vertically in each run. Run order was randomized. The initial object display was followed after 1000 ms by a centrally presented, informative arrow cue (60% valid) pointing to one rectangle end. The cue was followed after 1000 – 4000 ms by the target and distractors. The target could be presented in the same location denoted by the cue (valid trial), at a different location than the one denoted by the cue, but on the same object (invalid same-object trial), or at a different location on a different object than the one denoted by the cue (invalid different-object trial). The targets appearing in both the invalid locations were equidistant from the valid location. Distractors appeared in the non-target locations. Participants indicated whether the target was an X or an O, as quickly and accurately as possible by pressing one of two horizontally aligned buttons with their right hand. The mapping between target and response was counterbalanced. Targets remained on the screen for 2000 ms or until a response was detected, at which point a blank screen appeared for the remaining time. All blocks were randomized. The next block began after an interval of 1000 – 4000 ms.

**Figure 1.**
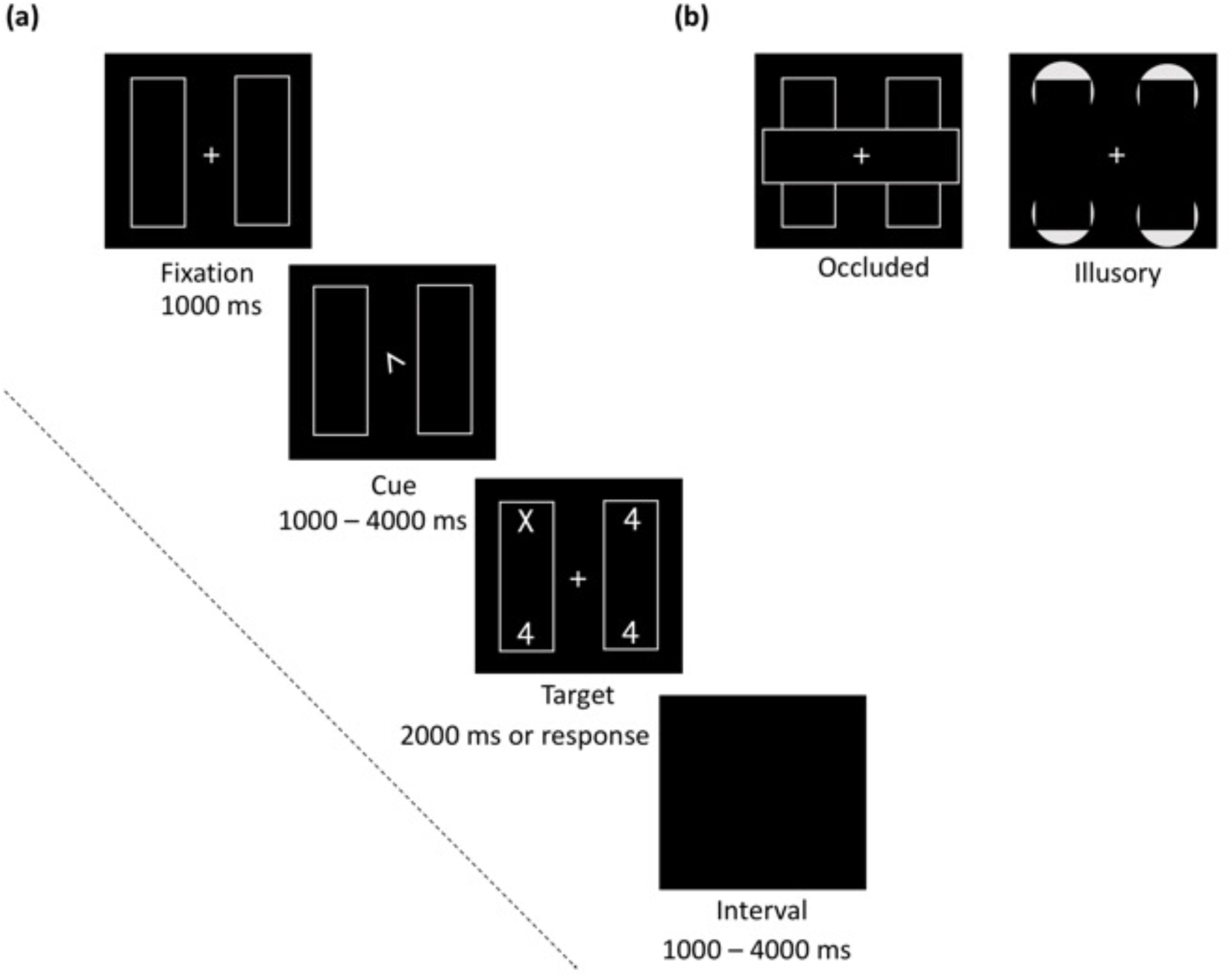
This figure represents the experimental design for the double-rectangle paradigm: **(a)** we depict a valid trial for explicit objects. The cue can point to any of the 4 possible target locations. Targets appeared in cued location on 60% of trials; and, **(b)** we show the occluded and illusory objects. All objects can be oriented horizontal or vertical.

#### fMRI procedure

All MR images were acquired using a 3 T GE scanner. Stimuli were back-projected onto a screen mounted to the scanner bore rear. Participants viewed the screen via a head coil-mounted mirror, and responded via an MR-compatible, fiber-optic two-button response box. We acquired T1-weighted spoiled gradient recalled acquisition in the steady state sequence volumes [repetition time (TR): 8.1 s; echo time (TE): 3.2 s; flip angle: 12^°^; 154 axial slices; voxel size: 1 mm^3^]. Moreover, we measured blood oxygenation level-dependent (BOLD) fMRI signal using a T2*-weighted EPI sequence [TR: 2.5 s; TE: 22 s; flip angle: 84^°^; 40 oblique slices; voxel size 2.7 mm^3^].

### Localizer Tasks

Along with the experimental task, we conducted 3 independent localizer tasks. These localizers were conducted across the 2 scan sessions, and comprised of a: **(1)** Meridian localizer scan to delineate the borders between V1, V2 and V3 in visual cortex; **(2)** Location localizer to identify patches in each region of visual cortex that respond to the 4 specific target locations (e.g., retinotopic area of cortex responds to the upper left location); and, **(3)** Lateral Occipital Cortex (LOC) localizer to identify object-selective regions of cortex. The procedures and stimuli for the 3 localizer tasks are detailed below:

#### Meridian mapping

Each participant performed 2 runs (309 s each) of the retinotopic meridian-mapping experiment, similar to previous studies (Greenberg et al., 2012; Uyar et al., 2016; see Fig. 2). Briefly, in this experiment, the borders between visual areas (dorsal and ventral V1, V2 and V3) were mapped by 8Hz checkerboard stimulation along the horizontal and vertical meridians. The initial fixation duration was 12 s, each meridian was presented – 8 times - for 18 s, and final fixation duration was 9 s. Participants were asked to indicate the change in color of a centrally presented square.

**Figure 2.**
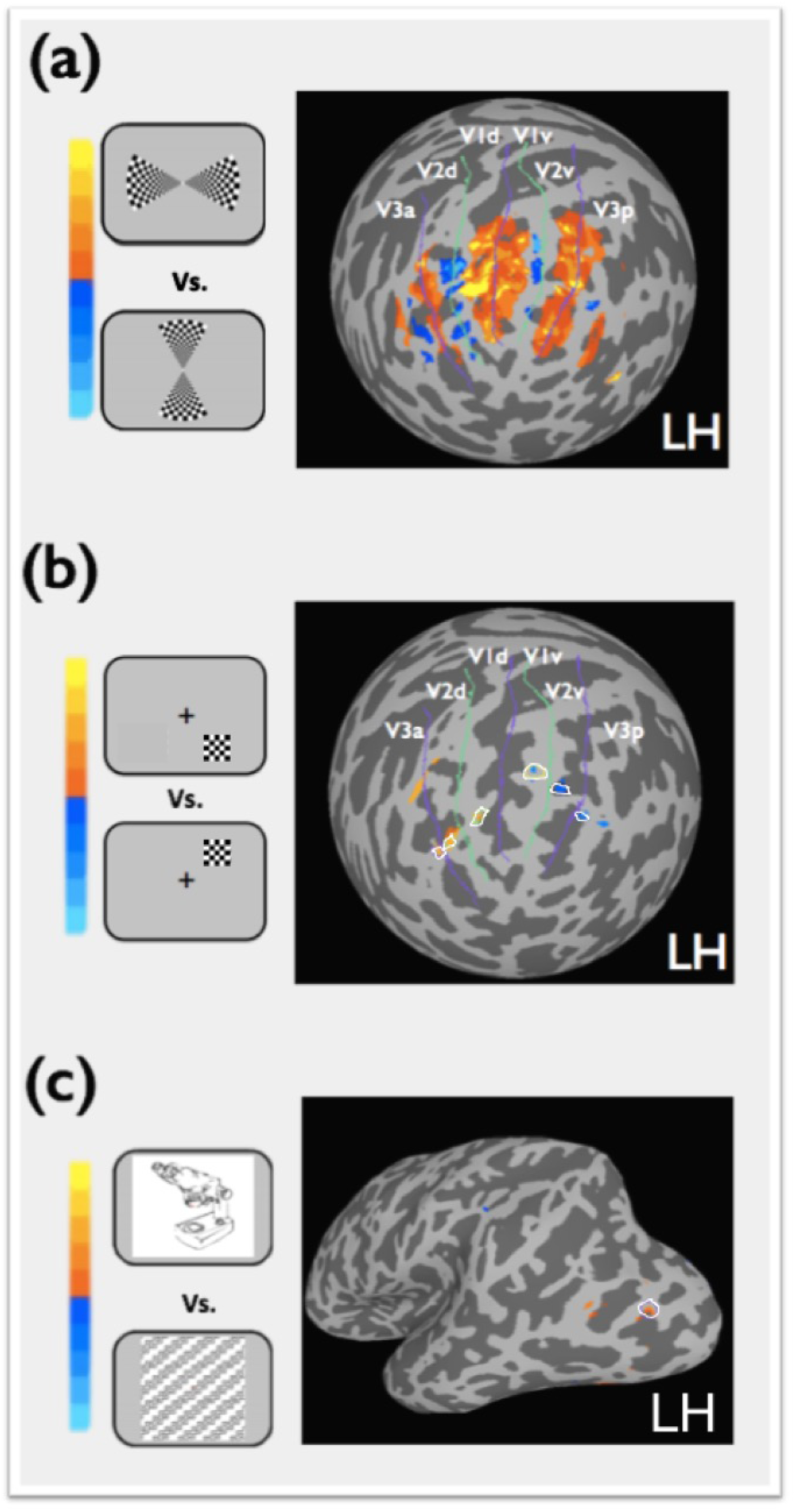
Shown here are the localizer tasks: **(a)** meridian mapping, which was used to delineate the (dorsal and ventral) V1 – V3 borders as revealed through a contrast of horizontal versus vertical meridian stimulation; **(b)** spatial localizer, which was used to identify the retinotopic representations of the 4-possible target locations in our experimental task – along the visual cortical hierarchy - as revealed through a contrast of lower right versus upper right stimulation for the left hemisphere, and upper left versus lower left stimulation for the right hemisphere; and, **(c)** LOC localizer, which was used to locate object-selective cortex as revealed through a contrast of images of objects versus patterns. Depicted here is the result of the contrasts, for each localizer, on an inflated left hemisphere brain (spherical and surface, respectively) for a representative participant.

#### Location localizer

Participants completed 2 runs (386 s) of the location localizer task, which is detailed in Uyar et al. (2016). For our purposes, a square of contrast-reversing checkerboards (at 4 Hz alternations) was presented. The square measured 1.36^°^ of visual angle, and was presented in the same coordinates as the edges of the rectangles used in the experimental task; that is, the locus of the targets in the experimental task. There were 20 blocks in total, 10 of each dyad (see Fig. 2). Initial fixation was 10 s and each block lasted 10 s, with 9 s fixation between each block and a final fixation of 5 s. Participants were asked to indicate the change in color of a centrally presented fixation.

#### LOC localizer

For the purposes of identifying regions of object-selective cortex, we asked participants to complete 2 runs (240 s) of an LOC localizer. The task included only blocks of objects and patterns, which were presented centrally (for details see Uyar et al., 2016; Fig. 2). Here, a stream of 9 stimuli were presented sequentially in a 9 s block. The initial fixation was 27 s, the interblock fixation was 6 s, and final fixation was 9 s. Participants completed a 1-back task; that is, indicate when an image appeared in a row.

### Statistical analyses

#### Behavioral data

Mean response latencies for correct responses was entered into a three-way ANOVA with Object Type (visible, occluded, illusory), Object Orientation (horizontal, vertical) and Cue-Target Validity (valid, invalid same, invalid different) as factors. The ANOVA was conducted using the ‘ez’ package in R (R Development Core Team, 2017). R was also used to calculate the mean accuracy across all our conditions.

#### Cortical surface reconstruction

The T1-weighted dataset was used to segment gray from white matter, and generate cortical surface representations for each hemisphere using FreeSurfer (e.g., Dale et al., 1999; Fischl et al., 1999; Greenberg et al., 2012).

#### fMRI preprocessing

Functional data were analyzed with the AFNI/SUMA software package (Cox, 1996; Saad et al., 2004), and custom MATLAB scripts. Before analysis, the first 4 volumes of each functional run were discarded. The remaining runs were slice-time corrected, and motion-corrected to the functional volume from the first run acquired in Session One. Data were then co-registered to the anatomical volume from which the surfaces were generated. The functional volumes from each run were next mapped to the cortical surface representation. The functional data was then converted to percentage signal change values normalized to the mean of each run, and spatial smoothing was performed. Statistical analyses were conducted on surface-mapped data.

#### Meridian mapping and regions of interest

There were a total of 7 ROIs in each hemisphere, for each participant. Six of those ROIs were from early visual cortex (dorsal and ventral V1, V2 and V3); hence, we first sought to delineate the visual cortex borders using the meridian localizer. We contrasted regressors from horizontal and vertical meridian conditions, and the borders between visual areas were then hand-drawn on the cortical surface by following the path of maximal activation, anteriorly from the occipital pole. Next, to determine ROIs that correspond to the retinotopic locations of the target (upper left, upper right, lower left, lower right), we used the location localizer. We contrasted regressors from upper left and lower left conditions in the right hemisphere, and upper right and lower right conditions in the left hemisphere. A 3mm ROI was grown from the resulting point of maximal activation within dorsal and ventral V1 – V3 using the ROIgrow function in AFNI. These ROIs allowed us to isolate cued, uncued same object and uncued different object locations for our analysis. Lastly, our seventh ROI was LOC, which was identified using the LOC localizer by contrasting object and pattern conditions. A 5mm ROI was grown from the point of maximal activation. Stimuli in the LOC localizer were centrally presented, hence the LOC is independent of visual field.

#### fMRI data

Event-related time courses in response to the cue were extracted from each ROI within retinotopic regions V1-V3, when that ROI was the currently attended position (i.e., cued location), the currently unattended position on the attended object (i.e., uncued same object location), and the currently unattended position on the unattended object (i.e., uncued different object location). These time courses were extracted separately for each object orientation, object type, location (dorsal/ventral) and hemisphere. For example, for the upper left location on a horizontal object (ventral V1-3, right hemisphere), only trials in which the cue directed attention to that locus, the same object locus (upper right), or the different object locus (lower left) were included, thus yielding cued, uncued same and different object locations, respectively. This analysis, thus, enabled us to investigate the activation pattern for an identical region in visual cortex (e.g., ventral V1-3, right hemisphere) when it served as the focus of the different attentional conditions. We then extracted the local maxima (first peak) in each of our resulting time courses using the ‘quantmod’ package in R (R Development Core Team, 2017). The peak values were submitted to a repeated-measures ANOVA with Region (V1, V2, V3), Object Type (visible, occluded, illusory), Object Orientation (horizontal, vertical), and Cueing (cued location, uncued same object location, uncued different object location). We discarded trials in which no peak was found for a given condition (0.03%).

We also extracted event-related time courses in response to the target from each ROI within retinotopic regions V1-V3. Here, we not only factored in whether or not that ROI was cued, but also whether or not the target then appeared at that ROI. We extracted, for each ROI, when attention was to be held at that location (hold trials), when attention shifted to it from another location on the same object (within-object shift), and when attention shifted to it from a different object (between-object shift). The time courses were extracted separately for each object orientation, object type, location (dorsal/ventral) and hemisphere. For example, for the upper left location on a horizontal object (ventral V1-3, right hemisphere), only trials in which the target appeared at that locus, and the cue directed attention to that locus, the same object locus (upper right), or the different object locus (lower left) were included, thus yielding hold, within-object and between-object shift conditions, respectively. We then extracted the local maxima. The peak values were entered into an ANOVA with Region (V1, V2, V3), Object Type (visible, occluded, illusory), Object Orientation (horizontal, vertical), and Event (hold, within-object shift, between-object shift). We discarded trials in which no peak was found (0.04%).

Lastly, we extracted event-related time courses in response to the target from LOC. Here, since LOC is not retinotopically-specific, we only factored in Cue-Target Validity. Essentially, these are the same conditions we used to extract the time series data from V1-V3. The time courses were extracted separately for each object orientation, object type, and hemisphere. We then extracted the local maxima in the resulting time courses. The peaks were entered to a repeated-measures ANOVA with the factors Object Type (visible, occluded, illusory), Object Orientation (horizontal, vertical), and Cue-Target Validity (valid, invalid same object location, invalid different object location).

## Results

### Behavioral results

The average accuracy across the 2 sessions was 93% (*SD* = 9). Our ANOVA with response latencies for correct responses revealed a three-way interaction, *F*(4, 84) = 4.091, *p* = 0.004. We investigated this three-way interaction by conducting a follow-up ANOVA with Object Type and Cue-Target Validity as factors, separately for each Object Orientation. The ANOVA for the horizontal objects revealed a significant main effect of Validity (see Fig. 3), *F*(2, 42) = 19.60, *p* < 0.0001. Subsequently, we calculated difference scores for the cue-validity effect (i.e., invalid same response latencies minus valid response latencies), and the same-object advantage (i.e., invalid different response latencies minus invalid same response latencies), when the objects were horizontal. We then conducted a one-sample t-test separately for each effect to examine whether or not the they were significantly different from zero. We found a 74 ms cue-validity effect, t(21) = 3.98, *p* = 0.0007. We also found a 22 ms same-object advantage, t(21) = 3.20, *p* = 0.004. In contrast, the ANOVA for vertical objects revealed a two-way interaction (Fig. 3), *F*(4, 84) = 2.56, *p* = 0.04. We found, here, a cue-validity effect for all object types (55 ms for visible objects, 102 ms for occluded objects and 88 ms for illusory objects; all *p*s < 0.01), but a same-object advantage of 29 ms emerged only for the visible objects, t(21) = 2.23, *p* = 0.04 (for inconsistent findings see Al-Janabi & Greenberg, 2016). All other t-tests were not significant (all *p*s > 0.05, and all *t*s < 1.90). Thus, we again find that object-based selection emergence is modulated by object orientation (Al-Janabi & Greenberg, 2016), such that the deployment of object-based attention is, at times, more efficient along the horizontal than the vertical meridian.

**Figure 3.**
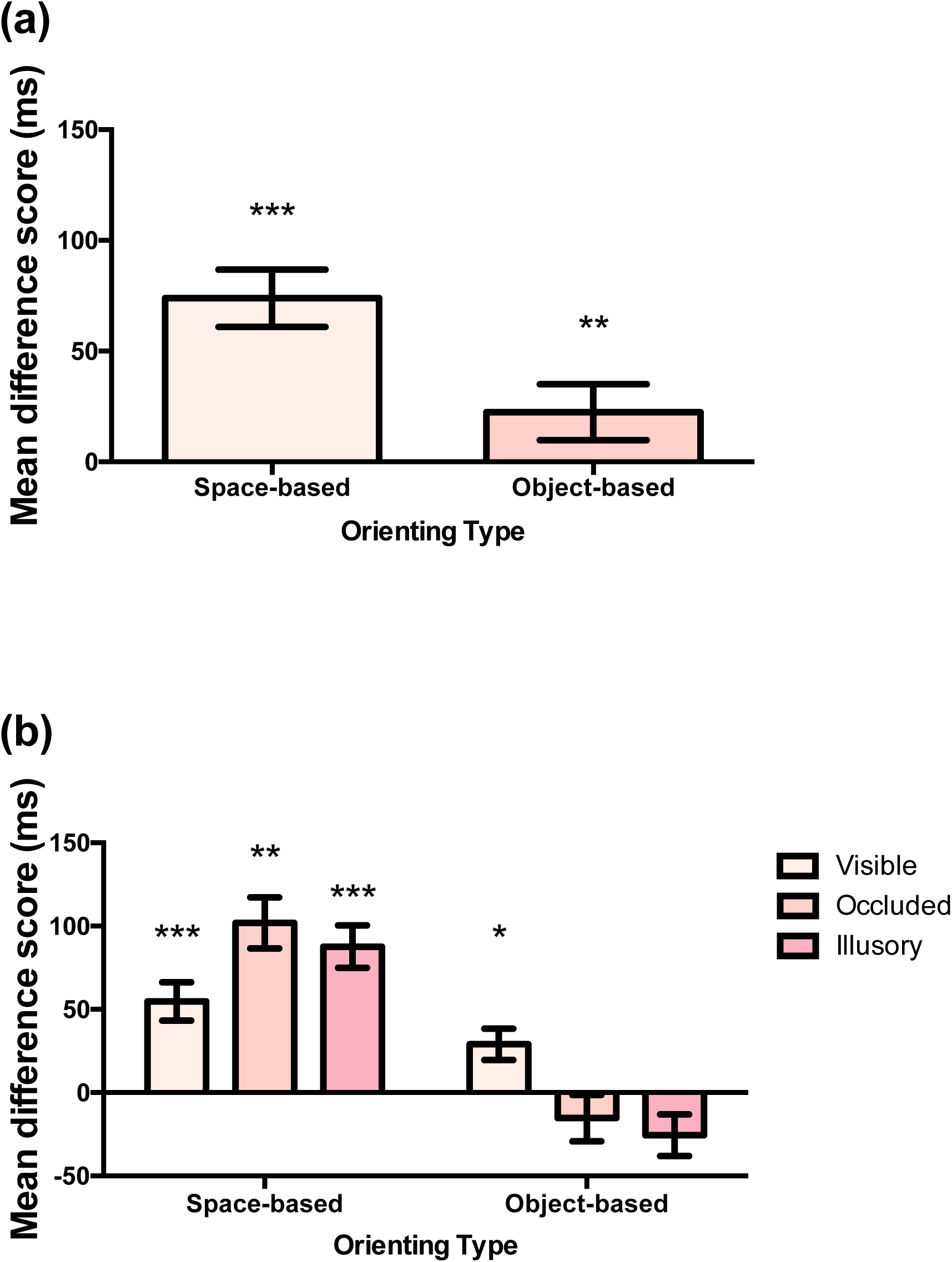
Here we show the mean difference scores for target identification when the objects are: **(a)** horizontal and **(b)** vertical. The mean difference scores (ms) for the cue-validity effect (space-based orienting) was calculated by subtracting valid trials from invalid same object trials. A positive score would be indicative of the effect. In contrast, the mean difference scores for the same-object advantage (object-based orienting) was calculated by subtracting invalid same from different object trials. A positive score would be indicative of the advantage. Our analysis indicates that neither effects were affected by Object Type when objects were horizontal. Error bars represent the within-subject standard error of the mean. Asterisks indicate * *p <* 0.05, ** *p* < 0.01, and *** *p* < 0.001.

### Cue-related activity results

Our omnibus ANOVA revealed no interactions between Object Orientation and Cueing (all *p*s > 0.05, *Fs* < 2.11), thus we collapsed across this factor. Our next ANOVA revealed two interactions with Cueing: one with Region, *F* (4, 84) = 3.20, *p* = 0.01, and Object Type, *F* (4, 84) = 18.57, *p* < 0.0001. We, therefore, conducted planned comparisons to investigate the nature of these interactions.

### Cueing and Region

We calculated scores that index cued location prioritization (i.e., cued location local maxima minus uncued same object location local maxima), and cued object prioritization (i.e., uncued different object location local maxima minus uncued same object location local maxima), separately for each visual area. We then conducted a one-sample t-test separately for each effect – in each area - to examine whether or not they were significantly different from zero. In all areas examined (see Fig. 4), we found no evidence of cued location prioritization (all *p*s > 0.05, and all *t*s < 1.80). Critically, however, we found evidence of cued object prioritization, but only in V3, t(21) = 2.36, *p* = 0.03. This result did not emerge in V1 or V2 (all *p*s > 0.05, and all *t*s < 1.90). It is important to mention, however, that the effect in V3 as compared to V1 and V2 was not significantly different (all *p*s > 0.05, all *t*s < 0.70). Our results, thus, indicate that BOLD signal enhancement following the cue was strongest on the cued object, and did not differ between locations on it. This latter result may reflect similar engagement of all target locations bound within a cued object.

**Figure 4.**
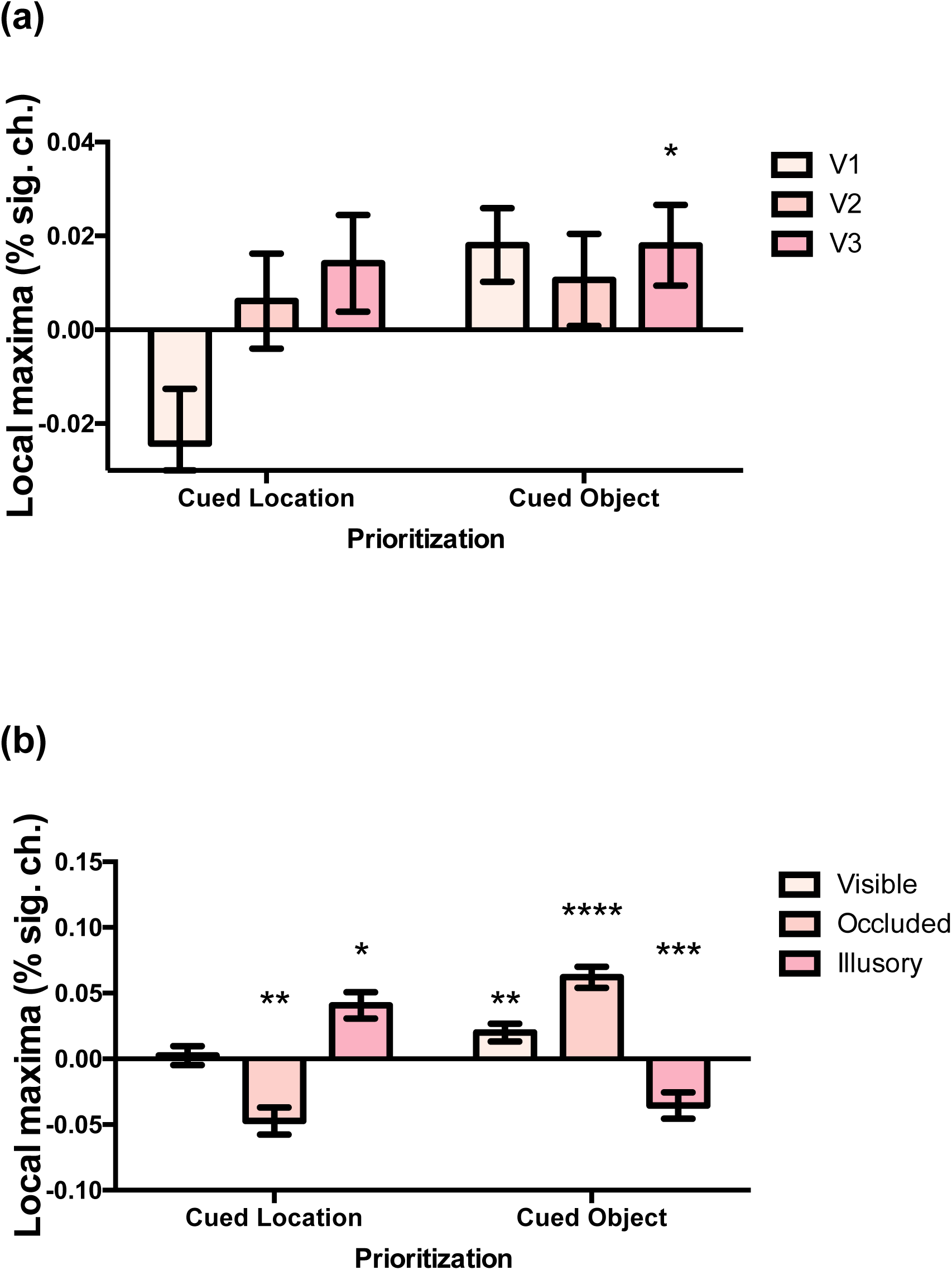
This figure depicts – following cue onset – prioritization of the cued location and cued object: **(a)** along the visual cortical hierarchy and **(b)** for each Object Type. Prioritization of the cued location was calculated by subtracting activation in retinotopic representations of the uncued same object location from the cued location. A positive score would be indicative of cued location prioritization. In contrast, prioritization of the cued object was calculated by subtracting activation in retinotopic representations of the uncued different from same object locations. A positive score would be indicative of cued object prioritization. Error bars represent the within-subject standard error of the mean. Asterisks indicate * *p <* 0.05, ** *p* < 0.01, *** *p* < 0.001, and **** *p* < 0.0001.

### Cueing and Grouping

As above, we calculated scores that index cued location prioritization and cued object prioritization, this time separately for each object type. We found that the emergence of cued location prioritization differed amongst the objects (Fig. 4), such that: **(a)** there was no evidence of such prioritization when the object was visible, t(21) = 0.23, *p* = 0.82; **(b)** there was a reversal of that prioritization for occluded objects, t(21) = 2.83, *p* = 0.01; and, **(c)** there was evidence of such prioritization for the illusory objects, t(21) = 2.78, *p* = 0.01. Similarly, the emergence of cued object prioritization also differed amongst the objects, such that: **(a)** there was evidence of such prioritization when the objects were visible, t(21) = 3.01, *p* = 0.007, or occluded, t(21) = 5.11, *p* < 0.0001; and, **(b)** there was no evidence of such prioritization when the objects were illusory, t (21) = 4.52, *p* = 0.0002. These results indicate that – following cue onset - object representation modulates cued location/object engagement^1^.

### Target-related activity results

Our omnibus ANOVA revealed no effect of Object Orientation on Event, *F* (2, 42) = 1.85, *p* = 0.17, thus we collapsed across this factor. Our subsequent ANOVA revealed two interactions with Event: one with Region, *F*(4, 84) = 2.66, *p* = 0.04, and one with Object Type, *F*(4, 84) = 2.73, *p* = 0.04. We, therefore, conducted planned comparisons to investigate the nature of these two-way interactions.

### Event and Region

We calculated scores that index between-location orienting (i.e., within object shifts local maxima minus hold local maxima), and between-object orienting (i.e., between object shifts local maxima minus within object shifts local maxima), separately for each visual area. We then conducted a one-sample t-test separately for each effect – in each area - to examine whether or not they were significantly different from zero. In all visual areas examined (see Fig. 5), we found evidence of between-location orienting (all *p*s < 0.01, and all *t*s > 2.80). Moreover, we found evidence of between-object orienting, but only in V3, t(21) = 2.09, *p* = 0.05. This result did not emerge in V1 or V2 (all *p*s > 0.05, and all *t*s < 0.50). It is important to mention, however, that the effect in V3 as compared to the effect in V2 was not significantly different, t(21) = 1.36, *p* = 0.19. Planned comparisons, thus, indicate that BOLD signal enhancement was strong when attentional shifts occurred between locations, as well as objects. This result may suggest target-related attentional reorienting from the cue location^2^.

**Figure 5.**
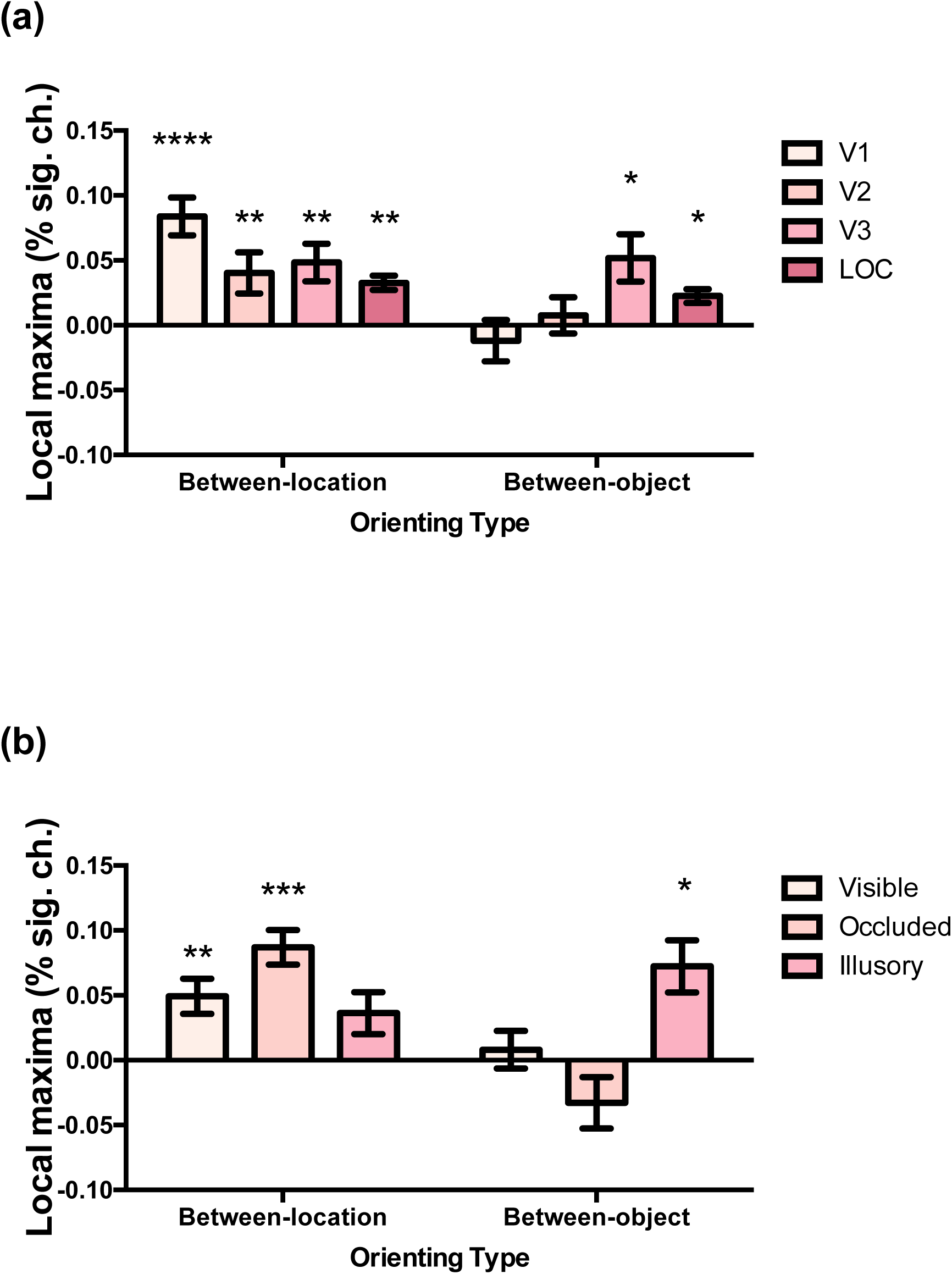
Here we show – following target onset – between-location and between-object orienting: **(a)** along the visual cortical hierarchy and **(b)** for each Object Type. Between-location orienting was calculated by subtracting activation in retinotopic representations of where attention was held from where it shifted within an object. A positive score would be indicative of between-location orienting. In contrast, between-object orienting was calculated by subtracting activation in retinotopic representations of where attention shifted within an object from where it shifted between objects. A positive score would be indicative of between-object orienting. Error bars represent the within-subject standard error of the mean. Asterisks indicate * *p <* 0.05, ** *p* < 0.01, *** *p* < 0.001, and **** *p* < 0.0001.

### Event and Grouping

As above, we calculated scores that index between-location and between-object orienting, this time separately for each object type. We found that the emergence of between-location orienting differed amongst the objects (Fig. 5), such that: **(a)** there was evidence of such orienting, but only for visible, t(21) = 2.90, *p* = 0.009, and occluded, t(21) = 4.57, *p* = 0.0002, objects; and, **(b)** there was no evidence of such orienting for illusory objects, t(21) = 1.33, *p* = 0.20. It is important to mention, however, that the between-location orienting effect for illusory objects as compared to visible and occluded objects was not significantly different (all *p*s > 0.05, all *t*s < 1.60). Similarly, we also found that the emergence of between-object orienting differed amongst the objects (Fig. 5), such that: **(a)** there was no evidence of such orienting for visible, t(21) = 0.32, *p* = 0.75, and occluded, t(21) = 1.12, *p* = 0.28, objects; and, in contrast, **(b)** there was evidence of such orienting for illusory objects, t(21) = 2.51, *p* = 0.02. It is important to mention, however, that the between-object effect for illusory objects as compared to visible objects was not significantly different, t (21) = 1.62, *p* = 0.12. These results indicate that objectness affects orienting to the target location^3^.

### LOC

Recall, for the target-related activity, we conducted a separate ANOVA to examine activation to our 3 attentional conditions in LOC (Fig. 5). This ANOVA revealed a significant main effect of Validity, *F*(2, 42) = 14.35, *p* < 0.0001. Specifically, we found evidence of both a cue-validity effect, t(21) = 3.52, *p* = 0.001, and same-object advantage, t(21) = 2.65, *p* = 0.01. These effects indicate that the between-location orienting effect we found in V1 – V3, and the between-object orienting effect we found in V3, are present in higher visual areas.

## General Discussion

The effects of object-based attention are not only limited to explicit object representations, but extend to object representations that require completion (Moore, Yantis & Vaughan, 1998). Our behavioral findings here support such a pattern of results. These results are intriguing because they imply that the processes underlying completion begin before object-based attention performs its function. One test of this implication is to investigate whether or not object-based effects for explicit object representations versus those that require completion emerge at the same level of visual cortex. Our study sought to carry out that test. Note, however, we did not intend to extrapolate from our findings when/where in visual cortex completion is accomplished, and attention then acts on the formed representation (for investigations on completion see Lerner, Hendler & Malach, 2002; Seghier & Vuilleumier, 2006), as that is beyond the scope of the coarse time-course resolution of imaging methodology. Rather, we sought to examine whether or not attention acts on object representations that do/do not require completion at the same level of visual cortex, which may point to differences in underlying mechanisms.

We observed several important findings in this study with regards to the initial orienting of attention in space and to objects following cue onset. First, we found that activity within retinotopic representations of target locations on a cued object increase to a similar extent along visual cortex. This result is consistent with the automatic spread of attention account (Vecera & Farah, 1994), which proposes that attention automatically and evenly spreads to all locations within the contour of an object that is signaled as relevant by a cue. In turn, the representation of the attended relative to the unattended object is enhanced. Second, and further confirming this cued object enhancement suggestion, we found that activity within retinotopic representations of the uncued same than different object locations is increased. This latter finding, though evident along visual cortex, was strongest in V3. Our findings from the cue period thus suggest that object-selection - when it is instantiated by a cue in the double-rectangle paradigm - may be started by attention similarly spreading to locations bound within a cued object. Our conclusion here is not dissimilar to that of Müller and Kleinschmidt (2003), with the exception that they, unlike us, show a graded spread of attention. The natural question is: for what reason does this discrepancy arise? To address this issue, we note that participants in the Müller and colleagues (2003) study always knew the cue location (upper left). Resultantly, it is likely that they used this information to pre-allocate attention to each target location prior to cue onset (during object exposure). This pre-allocation of attention may be graded because it was known that on 85% of trials the target will appear on the cued object: 70% of the time it will appear in the upper left location and 15% of the time it will appear in the upper right (for horizontal objects) or lower left (for vertical objects) same object locations. Thus, a priori knowledge of the cue location may have encouraged participants to split attention serially (Shomstein, 2012).

Our study also revealed several important findings about the reorienting of attention in space and to objects following target onset. First, we found that activity corresponding to retinotopic representations of where attention shifted within an object increased compared to where it was held. This between-location reorienting effect emerged along visual cortex, including LOC. Second, we found that activity corresponding to retinotopic representations of where attention shifted between versus within objects also increased. This between-object reorienting effect was strongest later in visual cortex (V3 and LOC), consistent with the biased competition account of attention (Bles et al., 2006)^4^. To our knowledge, this study is the first to show the reorienting of object-based attention in visual cortex, since neither Müller and Kleinschmidt (2003) nor Shomstein and Behrmann (2006) measured activity in retinotopic representations of the un/cued locations when the target appeared at those loci. Of course, our finding that activity increases when attention reorients between objects, and, to a lesser extent, locations begs the question: what does this activity increase reflect? There are two possible accounts. The extent to which activity increases when attention is reorienting in space or to objects may index **(1)** changes in alertness when a target appears at an unexpected location or object (Corbetta et al., 2002); or, **(2)** the effort associated with disengaging from the cued location and engaging the target location (on either the same or different object). This information may be received by visual cortex from the right intraparietal sulcus, which may respond to violations of expectancy, or the right temporo-parietal junction, which may be involved in disengaging attention (Thiel, Zills & Fink, 2004). Regardless of what these activity increases index, we show that there is a positive correlation between target-related visual cortex activity and behaviour. Our finding of faster responses to targets appearing at cued, followed by uncued same then different object locations may have a neural basis in visual cortex.

Given our interest in investigating whether or not object-based effects for explicit object representations versus those that require completion emerge at the same level of the visual cortical hierarchy, it would be remiss of us to not note the following: the object-based effects we observed following both cue and target onset emerged at the same level of visual cortex for all object representations. This finding indicates that not only is the behavior produced by explicit objects and those that require completion indistinguishable, but there is also no difference in the underlying neural basis of object-selection – at the level of visual cortex. The lack of a difference amongst object representations at either the behavioral or neural level may be because completion begins before object-based attention is deployed (within ∼100 ms; Woke et al., 2013). Indeed, given that partial object occlusion is the norm in the natural scene, efficient object-based selection of completed objects is well motivated. But, it is important to mention that although a same-object advantage was produced for all object representations, it is possible that the neural strategy invoking this effect differs amongst object representations. We show that, following cue onset, the pattern of activity for visible and occluded objects more closely follows an even spread of attention, whereas illusory objects follow a graded spread of attention. We also show that, following target onset, reorienting attention between-locations may be more difficult/linked to greater changes in alertness for visible and occluded versus illusory objects. In contrast, we show reorienting attention between-objects may be more difficult/linked to greater changes in alertness for illusory versus visible and occluded objects. These results, simply, reflect that object-selection for our object percepts, which vary in strength of object-hood, manifest in different ways. This finding is not wholly surprising; Marino and Scholl (2005) have shown that object-based effects can be enhanced or weakened by multiple cues to object-hood. The precise suggestion here by Marino and Scholl (2005) is that cues to object-hood affect the flow (or spread) of attention. Investigating the strategy underlying object-based effects for explicit and completed objects is, therefore, an avenue for future work.

## CONCLUSION

This study uncovers the neural basis underlying object-specific orienting and re-orienting of attention for explicit objects and those that require completion. Our results implicate later areas of visual cortex (V3 and LOC) in this process, which further supports accounts indicating that biased competition has its effect in later stages of the visual cortical hierarchy. Of import, these results suggest that the mechanism contributing to the selection of information on the basis of objects is an automatic spread of attention. It is by virtue of such a mechanism in visual cortex that attentional resources are preferentially and similarly allocated to locations bound within the contour of a relevant object, whilst suppressing locations bound within the contour of an irrelevant object.

## Acknowledgments

The authors would like to thank Mrinmayi Kulkarni and Dick Dubbelde for help with data collection. This work was supported by US-Israel Binational Science Foundation grant number 2013400 (A.S.G. & S.G.), as well as the University of Wisconsin – Milwaukee Research Growth Initiative (A.S.G.).

## Footnotes

1 In an attempt to understand these effects, we investigated the correlation between cued location prioritization and cued object prioritization for each object type. We found that the strength of association is negative when objects are visible (r = −0.31, *p* = 0.18) and occluded (r = −0.5, *p* = 0.02). This finding reflects that, to a moderate extent, <u>weaker</u> engagement of (solely) the cued location is correlated with stronger engagement of the cued object, which is commensurate with the cue-related functional effects we find for visible and occluded objects. In contrast, we found that the strength of association is positive when objects are illusory (r = 0.29, *p* = 0.17). This finding suggests that, to a small extent, <u>greater</u> engagement of (solely) the cued location is correlated with greater engagement of the cued object. Although the correlation between cued location and cued object prioritization is significant for only the occluded objects, the difference in correlation direction across objects may indicate that the strategy (graded vs. equal spread) participants use to allocate attention, after cue onset, is modulated by object type.

2 We conducted an additional analysis to investigate whether or not target-related activity in visual cortex is associated with behavioral performance, based on the assumption that both variables may index difficulty of attentional re-orienting from the cue to the target (e.g., Muller et al., 2003). We found that the strength of association between the cue-validity effect and its equivalent in the functional data (between-location orienting) is positive in V1 (r = 0.49, *p* = 0.03), V2 (r = 0.19, *p* = 0.34) and V3 (r = 0.23, *p* = 0.27). These results indicate difficulty re-orienting to the target <u>location</u> (i.e., large cue-validity effect) is mirrored in the functional data (i.e., greater between-location orienting), particularly in V1. We also found that the strength of association between the same-object effect and its equivalent in the functional data (between-object orienting) is positive in V1 (r = 0.52, *p* = 0.01), V2 (r = 0.44, *p* = 0.04) and V3 (r = 0.43, *p* = 0.05). These results indicate that the extent to which participants have difficulty re-orienting to the target <u>object</u> (i.e., larger same-object advantage) is also mirrored in the functional data (i.e., greater between-object orienting). Hence, to some extent, there is a relationship between functional activity and behavioral performance following target onset, indicating that the target-related activity, like behavior, may capture orienting to the target location.

3 We fleshed out this effect in two ways. First, we examined whether or not this modulation of orienting by grouping emerges in the behavior. Planned comparisons revealed that a cue-validity effect emerges for the visible (*M* Δ = 69 ms), occluded (*M* Δ = 90 ms), and illusory (*M* Δ = 75 ms), objects (all *ps* < 0.01, all *t*s > 3.50). However, the illusory objects showed the smallest effect size (Cohen’s *d* = 0.39), as compared to the visible (Cohen’s *d* = 0.44) and occluded (Cohen’s *d* = 0.50) objects. This result is consistent with our finding that the magnitude of between-location orienting was numerically, though not significantly, smallest for the illusory versus visible and occluded objects. Planned comparisons also revealed that a numerical same-object advantage emerges for the visible (*M* Δ = 13 ms), occluded (*M* Δ = 7 ms), and illusory (*M* Δ = 8 ms) objects (all *p*s > 0.05, all *t*s < 1.40). Critically, the occluded objects showed the smallest effect size (Cohen’s *d* = 0.039), as compared to the visible (Cohen’s *d* = 0.068) and illusory (Cohen’s *d* = 0.042) objects. This result is consistent with our finding that the magnitude of between-object orienting was smallest for occluded versus visible and illusory objects. Recall, accounting for the small same-object advantage reported here is an interaction between orientation and validity in the behavioral, not functional, data. Nonetheless, these findings suggest that the effect of grouping is not specific to target-related activity, but is also evident in behavior. The consistency in the results further indicates that both measures may index orienting from the cued to the target location. Second, we examined the extent to which between-location and between-object orienting correlate. We found that the strength of association is negative for visible (r = −0.64, *p* = 0.002), occluded (r = −0.46, *p* = 0.03), and illusory (r = −0.65, *p* = 0.001) objects. This finding suggests that increased activity associated with shifting between-locations (as indicated by a greater between-location orienting index) is correlated with decreased activity associated with shifting between-object (as indicated by a smaller between-object orienting index), and vice versa. This result is commensurate with both the behavioral effects we detailed above, and the target-related effects we find for all object types. These correlations may indicate that the ease with which the cued location is disengaged affects the ease with which shifts between locations/objects occurs; that is, if it is **(a)** difficult to disengage from the cued location and engage the uncued same object location then <u>there is no</u> added difficulty associated with disengaging from the cued object and engaging the uncued object; and, **(b)** easy to disengage from the cued location and engage the uncued same object location then <u>there is</u> added difficulty associated with disengaging from the cued object and engaging the uncued object. The extent to which **(a)** and **(b)** emerge may depend on whether the cued location is prioritized above the same object location following cue onset, thus making it difficult to disengage from that location and engage other locations, versus target locations on a cued object are similarly prioritized following cue onset, thus making it difficult to disengage from that object and engage other objects.

4 The biased competition account posits that attention can bias competition for neural representation between multiple stimuli in favor of an attended stimulus. Relieving that attended stimulus of the suppressive influences of distractors emerges, however, only later in the visual cortical hierarchy because receptive field sizes are larger – the exact locus in visual cortex depends on the magnitude of spatial separation between stimuli.

